# Isolating the key factors defining the magnitude of hippocampal neurogenesis’ effects on anxiety, memory and pattern separation

**DOI:** 10.1101/563023

**Authors:** Thiago F. A. França

## Abstract

In this paper, I analyze the hypothesis that hippocampal neurogenesis (HN) exerts its effects on behavior via activation of inhibitory circuits in the hippocampus. Using a very simple mathematical model (half-borrowed from biochemistry) to aid the reasoning, I show that the key factors determining the magnitude of HN’s effects on behavior are: the baseline levels of HN in the animal, the efficiency of the animal’s inhibitory circuits, the strength/intensity of the stimulus presented to the animal and how much accuracy the behavioral task requires from the information contained in the hippocampal representations. Taken together, those factors can help explain patterns observed in the behavioral results for memory, pattern separation and anxiety. The conclusions of the analysis suggest that HN’s effects on inhibitory circuits can explain the impact of neurogenesis on both emotion and cognition and provide a framework to interpret future studies about the effects of HN on different behaviors, with animals of different ages and of different species.

## 1. Introduction

Hippocampal neurogenesis (HN) has been a focus of intense research in the past decades (see Gonçalves et al. (2016) for a general review). Much of this research effort focused on understanding HN’s functional impact on the workings of the hippocampus, with an emphasis on its effects on behavior. We now have behavioral evidence for HN’s involvement in anxiety (see the meta-analysis in Groves et al., 2013), memory (Groves et al., 2013; França et al., 2017) and pattern separation (França et al., 2017). However, the effects of HN on those behaviors are not always clear. Although researchers have picked up some patterns in the behavioral results (Cameron and Glover, 2014), there is still a lot of ground to cover in the way of tying all these pieces of evidence together.

A promising hypothesis for a mechanism behind HN’s effects on behavior proposes that immature neurons in the dentate gyrus (DG) can activate local inhibitory circuits, enhancing feedback inhibition. HN may also affect CA3, contributing to feedforward inhibition in that sub-region (Piatti et al., 2013). Most of the studies regarding HN’s effects on inhibitory circuits focused on the DG, where there is evidence of HN ablation increasing granule cell activity/excitability (Burghardt et al., 2012; Lacefield et al., 2012; Ikar et al., 2013; Park et al., 2015), including a direct demonstration of immature neurons’ capacity to activate inhibitory circuits (Drew at al., 2016). Importantly, as I will show below, this single mechanism of action has the potential to explain HN’s effects in emotion, memory and pattern separation.

My goal in this paper is to get a better idea of how the effect of HN on inhibitory circuits can account for the different behavioral effects of HN. To do so, I will try isolating the key factors determining the magnitude of the effect of HN on behavior. In the following paragraphs, we will develop a model for HN’s effects on the inhibitory circuits of DG and CA3; we will try to capture the basic features of the workings of inhibitory circuits in a piece of mathematical formalism that I will keep as simple as possible. Our goal will not be to make quantitative predictions or to build a “realistic” model of the hippocampus. What we will try to do is use mathematical formalism merely to aid our reasoning in identifying the key factors that influence HN’s effects.

## 2. The model

We will use a simple input-output equation with the form

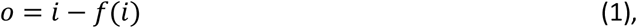

where *o* is the output activity of a given hippocampal sub-region (either DG or CA3) and *i* is the input to that region. The *f(i)* can stand for either feedback or feedforward inhibition, which in both cases is a function of the input. For now, we will focus on DG, with *f(i)* as feedback inhibition, but everything that will be developed below also applies to feedforward inhibition in CA3, as we shall see.

One caveat of using an input-output equation instead of, e.g., a computational model with “neurons” in it, is that we become blind to the identity of the neurons in the output, and this identity is of crucial importance for the workings of the hippocampus. However, for our purposes, we can circumvent this problem by organizing all the granule cells in the DG in a circular plane in a very particular way. To do so, consider a given stimulus to the DG, in the form of activation of several axons from the entorhinal cortex (EC) projecting to the DG. Given such stimulus, we will organize our DG granule cells in such a way that the neurons with the highest number of active synaptic inputs will be at the center of the circle, with neurons further away from the center having an increasingly smaller number of active inputs, all the way down to neurons with only one active synaptic input (we shall ignore neurons with no active inputs) (Figure 1a). In this context, the input *i* in our model will be the number of neurons receiving enough stimulation to, in the absence of any inhibition, reach the firing threshold. Thus, the further away from the center of the circle a neuron is, the smaller the number of active inputs it has and, consequently, the stronger the stimulus required to activate it. Therefore, the radius of the circle containing the granule cells activated by the input will be proportional to the strength or intensity of the stimulus – that is, how active where the EC neurons projecting to the DG granule cells. Having our circular arrangement of neurons in mind, the variable *i* will be defined as

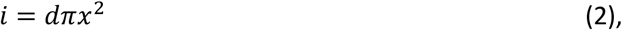

where *d* is the number of neurons per unit of area in our circular plane and *πx²* is the area of the circle containing the active neurons, the radius *x* being proportional to the intensity of the stimulus. We can see from the equation above that stronger stimuli will result in a larger value of x and, consequently, a larger circle encompassing a greater number of neurons. Notice that very high values of *x* would create a circle of active neurons so large that it would encompass even the neurons at the edge of our circular arrangement, representing the unnatural scenario in which stimulation is so strong, that even neurons with only one active synaptic input would receive enough neurotransmitter from that synapse to reach the firing threshold. The output *o* will be the number of neurons in the DG that actually become active after the effect of feedback inhibition. You can see now that, in the context of our model, the magnitude of the input is related to the intensity of our chosen stimulus, and an increase in the output for a given stimulus implies the activation of neurons with a smaller number of active synaptic inputs.

**Figure 1.**
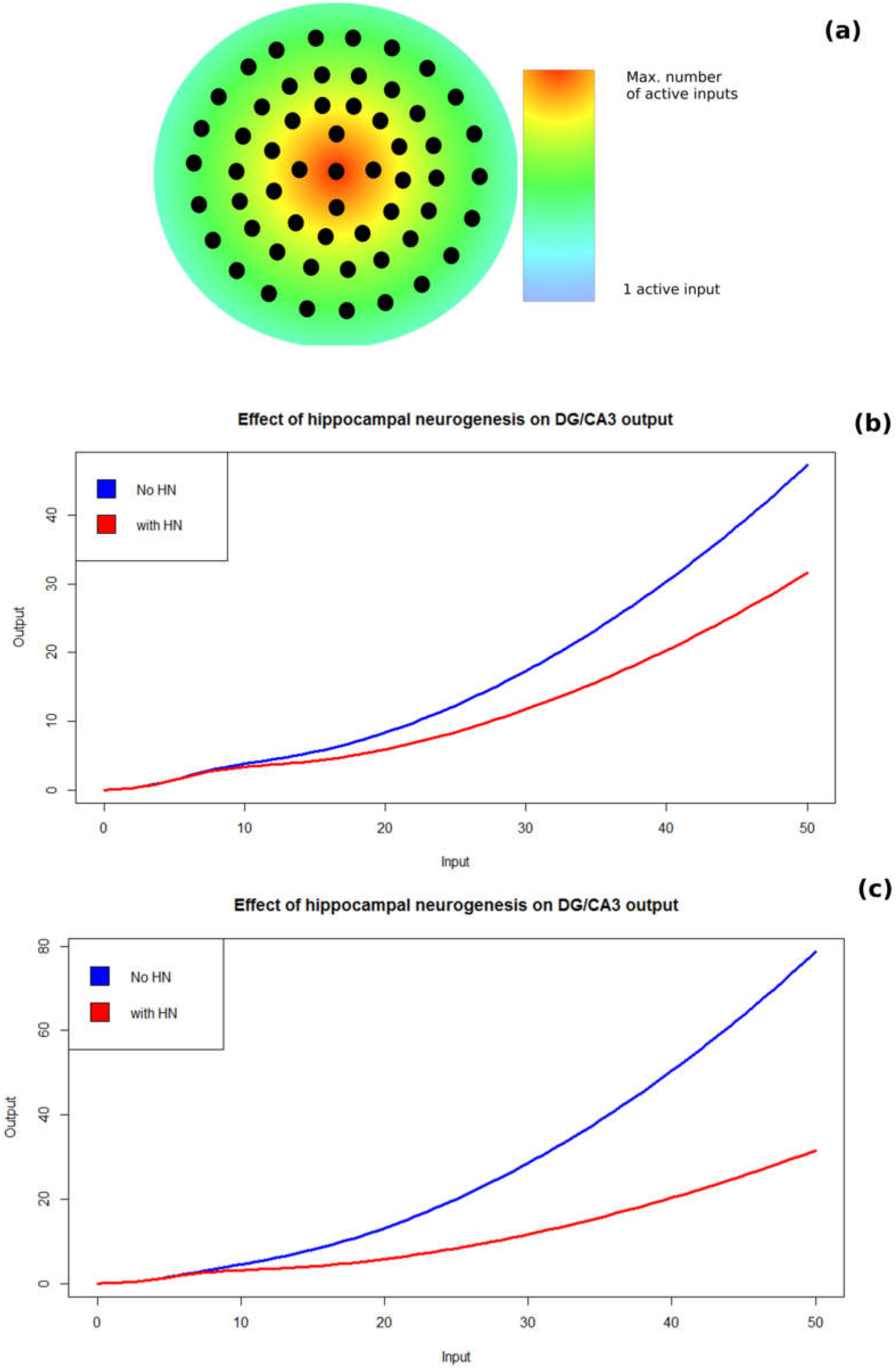
A simple model for hippocampal neurogenesis’ (HN) effect on the DG’s output via activation of feedback inhibition. **(a)** To simplify the mathematics of the model, we focus on a specific stimulus and arrange neurons (black dots) in a circular plane. The position of the neurons in the plane is defined by the number of the neurons’ synaptic inputs that are activated by the stimulus (note the color code). Thus, neurons with the highest number of synaptic inputs activated by the stimulus are at the center of the circle, while neurons with only one synaptic input activated are at the periphery. The radius of the area within the circle containing the neurons recruited by the input stimulus is proportional to the stimulus’ intensity. The role of feedback inhibition here is essentially to reduce the size of the area containing active neurons. **(b)** Comparison of equations 6 (blue curve) and 7 (red curve) for *K* = 30, *k* = 15, *c*=*h*=2, *d*=0.01, *M*=0.7 and *m*=0.1. The y axis represents the output and x axis represents the input. Note how the two curves diverge as the stimulus intensity, represented by the input’s magnitude, increases. **(c)** Same as (b), but with *M* = 0.5 and *m* = 0.3. Note how changing the parameters to give less weight to inhibition recruited by mature neurons and more weight to inhibition recruited by immature neurons accelerates the divergence of the curves.

We now come to the feedback term, *f(i)*. The strength of the feedback inhibition depends on the number of inhibitory neurons active, which depends on the intensity of the stimulus. Note that the whole process is teeming with cooperation since neurons combine their inputs over time and space. We can make an analogy with one of several molecular processes that involve cooperation. Take, for example, the binding of transcription regulators to regulatory regions of the DNA (Alberts at al., 2015). Such transcription regulators often act in dimers or even oligomers, needing to cooperate to elicit their effects, much like inhibitory interneurons must act together to efficiently inhibit principal neurons. Consider a situation with many transcription regulators and many DNA loci to bind. In such scenario, we would expect the relationship between transcription regulator concentration and their successful binding to DNA to be similar to the relation between the population activity of inhibitory interneurons and their inhibitory effect. We can thus borrow the Hill equation

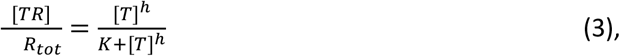

which describes the relationship between the concentration of transcription regulators [T] and the fraction of the total DNA loci that is bound to regulators ([TR]/R_tot_), with K being the dissociation constant and h being the Hill coefficient. What we are going to do is reinterpret the Hill equation for our purposes. We define

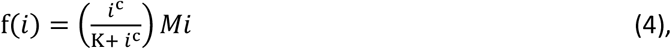

where we keep *i* = *dπx*^2^; *c* is the adapted Hill coefficient, which will be proportional to number of interneurons required to elicit a given inhibitory effect; *K*, the dissociation constant in the Hill equation, is the value of *i* required to elicit half of the feedback’s maximum capacity. The value of *K* will depend on, among other factors, the interneurons’ membrane time constant – a larger membrane time constant implies a small value of *K* and facilitates the summation of inputs by the neuron, just like a small dissociation rate facilitates DNA binding in the case of transcription regulators. Finally, *M* is the maximum inhibitory capacity – the maximum fraction of active principal neurons the feedback can inhibit, setting the maximum efficiency of feedback inhibition (note that the product *Mi* has no counterpart in the original Hill equation, it was added to convert the output of the Hill equation, which is a ratio, into a number of neurons to be subtracted from the input *i* in equation 1).

We know that neurons that project to the same targets cooperate to activate or inhibit their targets, and inhibitory interneurons show high conversion of inputs and divergence of outputs (McKenzie, 2018). But beyond the analogy between the Hill equation’s biochemical context and the feedback inhibition’s neural-network context, it is also worth noting that the shape of the Hill equation is a good match to observed input-output functions of hippocampal neurons (e.g., Pouille et al., 2013) – both show a sigmoidal pattern. In addition, the results reported by Drew et al (2016) on the evoked inhibitory potential in response to optogenetic stimulation of immature neurons also show a sigmoidal pattern after a very short initial period of almost vertical increase (see Figure 3g in Drew et al (2016)) However, one may object that, for feedback inhibition, there should be three nested Hill equations, one serving as input to the other, to represent the three stages where there is cooperativity: in the activation of principal neurons by the input stimulus, in the activation of inhibitory interneurons by principal neurons, and in the inhibition of principal neurons by interneurons. However, such equations, with the general form

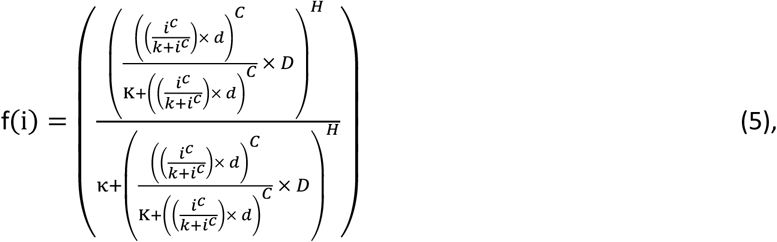

would produce curves that maintain the sigmoid shape characteristic of the Hill equation with cooperativity, as equation 5 maintains the same overall structure of the Hill equation ([T]^h^/K +[T]^h^). Since we are only interested in qualitative analysis here, we can go without using such an ugly equation.

We are now in position to plug all the terms (equations 2 and 4) in our original equation, *o* = *i* − *f*(*i*), to produce

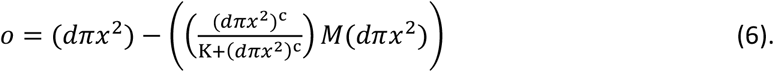

How do we represent HN in this equation? By adding a second feedback term, with the exact same form as the feedback term in equation 4:

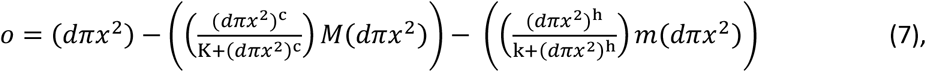

where *k, m* and *h* are defined as *K*, *M* and *c*, respectively, in the first feedback term (equation 4). We set the first feedback term to represent the feedback inhibition elicited by mature granule cells, while the second feedback term represents the feedback recruited by immature granule cells. Note that we are simplifying things by assuming that the two feedback terms add linearly. An alternative formulation would be to model the feedback elicited by both mature and immature neurons in just one feedback term, a Hill equation in which the maximum inhibitory capacity equals *M* + *m* and the “dissociation constant” falls between *K* and *k*. I show in the **Supplementary Figure 1** that this model shows the same behavior as the one we are using. I opted for this version with separate feedback terms for mature and immature granule cells because it will make it easier to visualize some points that we are going to discuss bellow.

Note that the equations above are also valid for feedforward inhibition in CA3. Like feedback inhibition in the DG, feedforward inhibition in and CA3 also shows cooperativity and can be described using the same equations by taking the input to be the stimulation of CA3 pyramidal cells by DG granule cells and the *f(i)* terms to be feedforward inhibition on CA3. Therefore, the conclusions we are about to draw can apply to HN’s effects both on DG and CA3.

## 3. Comparing the model with and without HN

What we are going to do now is to compare equations 6 and 7 to see how the output behaves with and without the HN feedback term. The specific choice of parameters does not really matter, as the same overall pattern can be observed for a wide range of parameters, as long as *M* + *m* < 1. Note that *M* + *m* > 1 would imply that stronger stimuli activate fewer neurons, as the maximum efficiency of the inhibition would be more than enough to inhibit every single neuron stimulated. We also assume that *M* > *m*, meaning that mature granule cells recruit more feedback/feedforwad inhibition than immature granule cells, although violating this assumption would imply a greater magnitude of HN’ effects without changing our conclusions as to the relevant factors influencing such effect. Finally, we also assume *K* > *k*, meaning that the inhibition recruited by immature neurons reaches the plateau faster since immature neurons are more excitable (Gonçalves et al., 2016).

Figure 1b shows the comparison of equations 6 (blue curve) and 7 (red curve) for *K* = 30, *k* = 15, *c*=*h*=2, *d*=0.01, *M*=0.7 and *m*=0.1 and Figure 1c shows the comparison after changing *M* to 0.5 and *m* to 0.3 (the interested reader can use the R code and the step-by-step guide provided in the **Supplementary Materials** to test other values and see that the general pattern observed for the example values is conserved). The main implications of the model can be visualized by analyzing the curves in Figures 1b and 1c. There are some key features that I would like to highlight.

### 3.1. Implications for HN’s effects on anxiety

The first feature I would like to highlight is that the model implies that the contribution of immature neurons to feedback/feedforward inhibition, which reflects in the difference between the outputs with and without HN, grows with the intensity of the stimulus – although it is worth noting that in a real situation, the output would eventually hit a plateau, as the number of granule cells and interneurons is finite. Importantly, this pattern agrees with experimental data regarding response strength of the DG in animals with increased or reduced HN compared to control (see Figures 2 and 3 of Ikar et al., 2013). This can explain the observation that a stronger impact of HN ablation on anxiety-related behaviors is obtained when animals are subjected to stress prior testing (see Cameron and Glover, 2014). Such observation fits well with our model, as more intense stimulation would make the effects of HN ablation more apparent. Accordingly, the expression of anxiety- and avoidance-related behaviors was shown to depend on the activity of neurons in the ventral hippocampus projecting to the lateral hypothalamic area (Jimenez et al., 2018), and a recent study strongly indicates that HN confers stress resilience precisely by inhibiting the ventral DG (Anacker et al., 2018).

### 3.2. Implications for HN’s effects on memory and pattern separation

From the context of our model, we can see that the surplus in the number of output neurons consists of neurons having an increasingly smaller number of active synaptic inputs (see our discussion of the model in Section 2 and Figure 1a). This has important implications for the cognitive effects of HN. Some authors have noted before that that the magnitude of neurogenesis’ effects on memory tasks seem to be greater in more difficult tests (Cameron and Glover, 2014). In the case of memory and pattern separation, this aspect of the model can give an explanation as to why the behavioral effect of HN seems greater and more consistent in studies employing behavioral tests of pattern separation compared to other memory tasks, such as Morris water maze and contextual fear conditioning (Groves et al., 2013; França et al., 2017). Note that the inhibitory circuits create competition between neurons by inhibiting the less active neurons and selecting the ones receiving stronger input (França and Monserrat, 2018). In doing so, inhibition helps to maintain active only the neurons whose receptive fields (as defined by their connectivity pattern) better represent the stimulus being given. The weakening of feedback/feedforward inhibition would ease the competition and allow the activation of neurons representing information that does not fit the stimulus so well, essentially adding noise to hippocampal representations (for a more detailed discussion of the effects of inhibition on hippocampal representations, see França and Monserrat (2018), Mckenzie (2018) and Rao-Ruiz et al (2019)). But how much would this added noise influence the animals’ behavior? As we have seen, that will depend on the intensity of the stimulus. But, importantly, it also depends on the difficulty of the test. The harder the test, the more detail an animal has to remember, the more likely it is that the animal will be impaired by the noise added by HN ablation. Since behavioral tests of pattern separation are the ones demanding the most of memory by presenting the animal with very similar stimuli, it would be expected that animals are more consistently impaired in such tests.

At this point, it is important to note that immature neurons, being highly plastic and excitable and firing with low specificity, could be themselves a source of noise in the DG’s output. However, synaptic transmission from granule cells to CA3 pyramidal cells is characterized by its high efficiency, and this efficiency stems from the peculiar architecture of those synapses. Such specialized structure takes very long to develop in new granule cells, with its development being completed only after the critical period of high excitability (Toni and Schinder, 2016). And while granule cells contact only a dozen or so CA3 pyramidal cells, filopodial extensions from mossy fiber terminals allows granule cells to contact more than twice as many GABAergic interneurons in CA3 (Acsády et al., 1998). Moreover, immature neurons have been reported to form more synapses with inhibitory interneurons in the CA3 compared to mature granule cells (Restivo et al., 2012). Taken together, these observations suggest that whatever is the direct contribution of immature granule cells to increase DG’s output, this contribution is offset by their indirect contribution to reduce DG’s output, as well as their inhibitory effect on CA3 and the delayed morphological maturation of their presynaptic terminals with CA3 pyramidal cells.

### 3.3. Individual differences, juvenile *vs.* adult HN, and implications for comparative studies

Another key implication of the model I would like to highlight is that the effect of HN ablation will depend not only on the contribution of immature neurons to feedback/feedforward inhibition, but also on the capacity of mature neurons to elicit feedback/feedforward inhibition. This can be seen by comparing Figures 1b and 1c. In both cases, the total inhibitory capacity (*M* + *m*) is 0.8; what determines the importance of HN for the animal is the way that total capacity is distributed. This could explain at least part of the variability observed in behavioral studies after HN ablation. Animals vary in their levels of HN at baseline and can also differ in the efficiency of their inhibitory circuits (which could be influenced by individual variations in, e.g., cell excitability due to the levels of neuromodulators/hormones or the connectivity pattern and synaptic weights between principal neurons and interneurons).

A particularly relevant example of individual variation occurs when comparing juvenile and adult animals. Not only are the rates of HN higher in juvenile animals compared to adults (Snyder, 2019), but there may also be changes in the efficiency of inhibitory circuits between juvenile and adult animals due to, for example, the increase in levels of sex hormones during puberty. Such hormones can alter the function of inhibitory interneurons the neocortex (Piekarski et al., 2017; Clemens et al., 2019) and likely do so in the hippocampus as well (Hart et al., 2001). Thus, HN may be more important for eliciting proper behavior in animals of specific age groups.

In the same token, HN may be more important for hippocampal function in some species than in others due to interspecies variations in feedforward/feedback circuits. This is a very important point when considering possible clinical applications of manipulating HN. While most studies of HN were conducting using rodents, there are significant differences between rodents and primates in numbers of inhibitory interneurons and in the connectivity pattern of inhibitory circuits (Freund and Buzsáki, 1996).

## 4. Testing against competing hypotheses

The model presented here shows that neurogenesis’ effects on inhibitory circuits *could* explain the evidence discussed, and I argued that the direct effect of immature neurons on CA3 pyramidal neurons is offset by their indirect effect via modulation of inhibitory circuits. However, we cannot completely rule out the possibility of immature neurons’ excitatory projections are affecting CA3 representations directly, and some researchers have even suggested that immature neurons can directly contribute information necessary for hippocampus’ proper functioning (e.g., Aimone et al., 2006; Aimone et al., 2011).

Experiments designed to specifically distinguish between direct and indirect effects of immature neurons are lacking. Such discrimination would require that we manipulate the two mechanisms separately. This could be achieved by combining optogenetic inhibition of immature neurons with optogenetic stimulation of inhibitory interneurons in the DG. Silencing immature neurons would block their direct effect on CA3 pyramidal cells while stimulating inhibitory interneurons would supply something equivalent to their indirect effect. If our model is correct, then silencing immature neurons would impair performance in behavioral tests of memory, pattern separation and anxiety, but simultaneously stimulating inhibitory interneurons would lead to a behavioral performance equal or better compared to control. Conversely, if behavioral effects of neurogenesis are mediated by immature neurons’ direct effect on CA3 representations, then the intervention described above would impair behavioral performance in all tests and stimulation of inhibitory interneurons would do little to improve the damage caused by silencing immature neurons. Finally, if both direct and indirect mechanisms are operating, then stimulating inhibitory interneurons would improve behavioral performance in animals with silenced immature neurons, but performance would not be comparable to that of control animals. In this way, future studies may be able to support or falsify our proposed framework.

## 5. Conclusions

In conclusion, we can see from the reasoning exposed above that the magnitude of the behavioral effect of HN, at least of the effects elicited via activation of inhibitory circuits, largely depends on just a few key factors. They include the levels of HN in the animal, the efficiency of the animal’s inhibitory circuits, the strength of the stimulus presented to the animal and how much accuracy is required from the information contained in hippocampal representations. Such principles may provide a coherent framework to interpret future results of behavioral studies in different species, whether they deal with cognition or emotion.

## Acknowledgements

The author would like to thank Dr. José Maria Monserrat for helpful comments and review of the manuscript. During the production of this work, the author received financial support from the Coordenação de Aperfeiçoamento de Pessoal de Nível Superior - Brasil (CAPES) – Finance Code 001.

