## supplementary material for "Isolating the key factors defining the magnitude of hippocampal neurogenesis’ effects on anxiety, memory and pattern separation"

In the following text I provide the codes used to plot the two model equations (equations 6 and 7 in the main text) using R. The text below provides a step-by-step guide to plot the function with different parameter values, so no prior knowledge of the software R is required; all you will need is to have R installed in your computer (R can be downloaded for free at https://cran.r-project.org/bin/) and paste the codes bellow (without the ">" signs) in the console window.

The first thing we need to do is put our equations in R. Both equations used in the model (see main text) are functions and can be used in R by writing:

```
>f = function(x){(d*2*3.14*(x^2))-(((((d*2*3.14*(x^2))^c)/(K+(d*2*3.14*(x^2))^c)))*(M*
(d*2*3.14*(x^2))))}
```

```
>g = function(x){(d*2*3.14*(x^2))-(((((d*2*3.14*(x^2))^c)/(K+(d*2*3.14*(x^2))^c)))*(M*
(d*2*3.14*(x^2))))-(((((d*2*3.14*(x^2))^c)/(k+(d*2*3.14*(x^2))^h)))*(m* (d*2*3.14*(x^2))))}
```

Here, f is equation 6 of the main text, without neurogenesis,

$$o = (d\pi x^2) - \left(\left(\frac{(d\pi x^2)^{\mathrm{c}}}{\mathrm{K} + (d\pi x^2)^{\mathrm{c}}}\right) M(d\pi x^2)\right),$$

and g is equation 7, with neurogenesis,

$$o = (d\pi x^2) - \left(\left(\frac{(d\pi x^2)^{\mathrm{c}}}{\mathrm{K}+(d\pi x^2)^{\mathrm{c}}}\right) M(n\pi x^2)\right) - \left(\left(\frac{(d\pi x^2)^{\mathrm{h}}}{\mathrm{k}+(d\pi x^2)^{\mathrm{h}}}\right) m(d\pi x^2)\right).$$

The next step is give values to the parameters. For the values used in Figure 1b of the main text, we write:

```
>d=0.01
>K=30
>k=15
>c=2
```

```
>h=2

>M=0.7

>m=0.1
```

For the values used in figure 1c, we write instead

```
>d=0.01

>K=30

>k=15

>c=2

>h=2

>M=0.5

>m=0.3
```

By changing the values in the text above, you can plot the functions with any parameter value you want.

Finally, to produce the plots, we use the plot() function and write:

```
p=plot(f, to=50,

    main="Effect of hippocampal neurogenesis on DG/CA3 output",

    ylab="Output",

    xlab= "Input",

    type="l",

    col="blue",

    lwd=3)

legend("topleft",
```

```
    c("No HN","with HN"),
    fill=c("blue","red")
)
plot(g, to=50, col="red",lwd=3, add=TRUE)
```

The code above plots both equations (regardless of the parameter values chosen) in the same graph and has arguments defining things like line color, axes names, plot title, etc. An important argument that you may like to change is the “to=50”, which tells R to plot the function all the way to x=50. By changing this value you can plot the functions for larger input values.
