## Supplementary Figure 1 for "Isolating the key factors defining the magnitude of hippocampal neurogenesis’ effects on anxiety, memory and pattern separation"

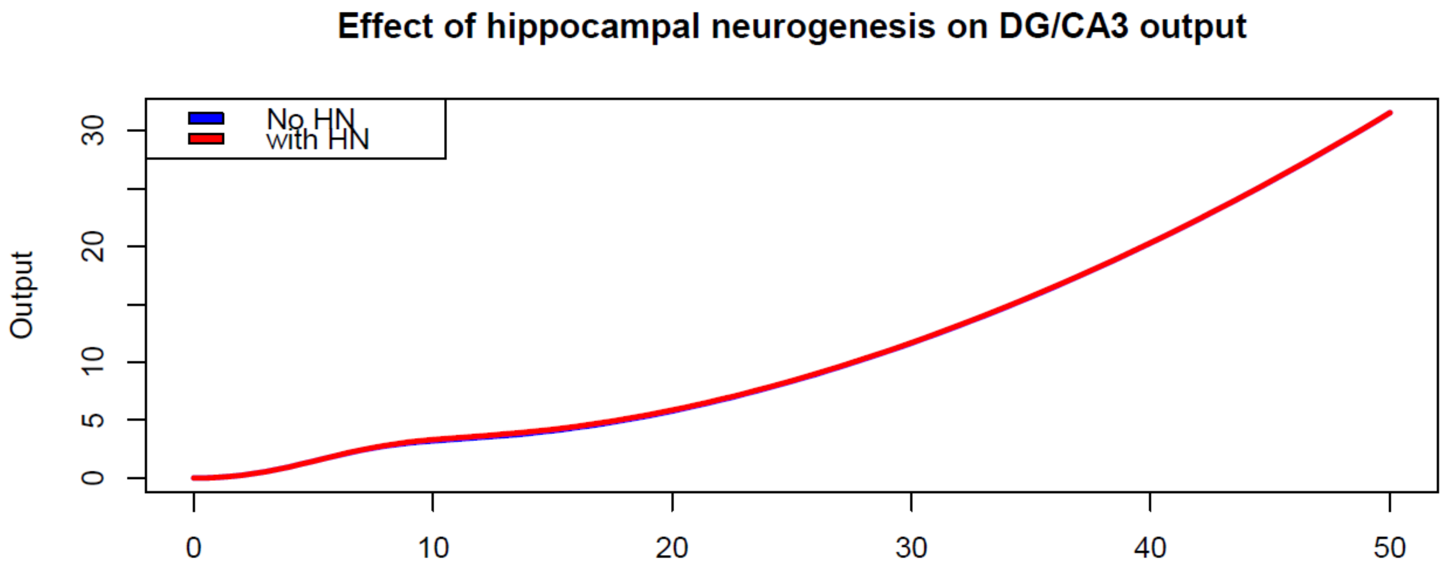


**An alternative way of representing HN in the model.** As mentioned in the main text, an alternative way to represent HN-elicited feedback in the input-output model is to incorporate it to the same feedback term that was representing mature granule cell-feedback. In this alternative version of the model, the maximum inhibitory capacity equals $M + m$ and the "dissociation constant" (the value of $i$ required to elicit half of the feedback's maximum capacity) is lowered to a value between $K$ and $k$. To show that those two formulations create the same results, I plotted equation 7 of the main text using the same parameters as in Figure 1a ($K$ = 30, $k$ = 15, $c$=$h$=2, $d$=0.01, $M$=0.7 and $m$=0.1) along with the alternative model, with $K$=22.5 (exactly halfway between K and K in Figure 1a), M=0.8 (which equals $M + m$ in Figure 1a) and the other parameters as in Figure 1a. The resulting plot shows that the two curves match perfectly.
